# CRISPR screening identifies novel PARP inhibitor classification based on distinct base excision repair pathway dependencies

**DOI:** 10.1101/2020.10.18.333070

**Authors:** Gregory A. Breuer, Jonathan Bezney, Nathan R. Fons, Ranjini K. Sundaram, Wanjuan Feng, Gaorav P. Gupta, Ranjit S. Bindra

## Abstract

DNA repair deficiencies have become an increasingly promising target for novel therapeutics within the realm of clinical oncology. Recently, several inhibitors of Poly(ADP-ribose) Polymerases (PARPs) have received approval for the treatment of cancers primarily with deleterious mutations in the homologous recombination (HR) proteins, BRCA1 and BRCA2. Despite numerous clinical trials which have been completed or are currently ongoing, the mechanism of action by which PARP inhibitors selectively kill tumor cells is poorly understood. While many believe “trapping” of PARP proteins to DNA at sites of damage is the most important determinant driving cytotoxicity by these drugs, clinically effective inhibitors exist with a diverse range of PARP-trapping qualities. These findings suggest that characterization of inhibitors as strong versus weak trappers does not properly capture the intra-class characteristics of these drugs. Here, we use a novel, targeted DNA damage response (DDR) CRISPR/Cas9 screening library to reveal heterogenous genetic dependencies on the base excision repair (BER) pathway for PARP inhibitors, which is not correlated with PARP trapping ability or catalytic inhibition of PARP. These findings demonstrate that inhibition of PARylation and induction of PARP trapping are not the only factors contributing to distinct biological activity for different PARP inhibitors, and they provide insight into the optimal choice of PARP inhibitors for use in the setting of specific DDR defects.

**AUTHOR SUMMARY:** Targeted cancer therapies rely on our general understanding of which genetic mutations are involved in both sensitivity and resistance to such anticancer agents. In this study, we describe the use of functional genetic screening to evaluate the role of various DNA repair proteins in response to inhibitors of PARP, a quintessential example of targeted therapy. While PARP inhibitors are best known for their utility in cancers with homologous recombination defects, we show that some inhibitors within this class may have additional functionality in cancers with deficient base excision repair. These findings highlight not only the importance of PARP inhibitor selection in the appropriate context, but also the mechanistic differences that exist within this class of inhibitors. It is our hope that our findings will inspire future work evaluating the use of specific PARP inhibitor selection in designing clinical trials to further expand the use of PARP inhibitors beyond tumors with homologous recombination deficiencies.

## INTRODUCTION

Over the last decade, inhibitors of poly(ADP-ribose) polymerase-1 and −2 (PARP1/2) have been established as safe and effective cancer therapeutics, which are most active against tumors with homologous recombination (HR) defects, such as those with deleterious mutations in BRCA1, BRCA2, and others [1–3]. The PARP family of proteins utilize NAD+ to add one (mono-) or more (poly-) ADP-ribose chains to target proteins in response to various stimuli (referred to as PARylation) [4]. While most proteins downstream from PARP1 and PARP2 act in DNA damage response (DDR) pathways, over 170 different PARP interactions have been described, and thus these proteins play important roles in a diverse range of functions, ranging from cell cycle regulation to cell motility [5, 6]. Furthermore, the targets of such PARylation events are known to be stimulus-dependent [7]. PARP proteins play a well-established role in single strand break (SSB) repair, in which they recruit proteins such as XRCC1 and other factors for resolution of these lesions [8]. Prevention of SSB repair can result in increased replication stress, unrepaired double-strand breaks (DSBs), and difficulty with replication restart, which collectively are thought to underlie the enhanced cytotoxicity of PARP inhibitors in HR-defective cancers [9–11]. However, recent evidence suggests that the effect of “trapping” PARP1 at sites of SSB repair may be more important for cytotoxicity of these agents, particularly in HR-defective cells [12, 13]. Trapping has been exhibited for both PARP1 and PARP2, though PARP1 remains the most important family member regarding SSB repair and the induction of synthetic lethality [8]. Despite these new insights, clinically relevant PARP inhibitors exist across a wide spectrum of potencies and specificities, in relation to PARP trapping ability, catalytic inhibition of PARylation, and efficacy in targeting other members of the PARP family of proteins [14]. Additionally, loss of PARP function in the setting of HR deficiencies shows moderate growth inhibition, independent of trapping inhibitors, indicating that both actions may be important for cell toxicity [15].

More recent studies suggest that synthetic lethal interactions with PARP inhibitors extend beyond BRCA1 and BRCA2 mutations, to including additional DDR proteins, such as mutations in the RAD51 paralogues, PALB2, ATM, and others [16]. Furthermore, PARP inhibitor sensitivity has been used as a screening tool to identify novel HR-related functions of genes, such as mutant IDH1 and ribonuclease H2 [17, 18]. As *in vitro* studies continue to show an ever-expanding landscape of possible uses for PARP inhibitors, it is not fully understood whether these sensitivities extend across the entire class of PARP inhibitors, or only a subset of drugs within this class.

With the knowledge that PARP trapping ability is functionally independent of catalytic inhibition, we set out to characterize the utility of the clinically available inhibitors -- olaparib, rucaparib, talazoparib, niraparib, and veliparib. Using a high coverage, targeted DDR CRISPR/Cas9-based screening library, we have developed a novel assay focused solely on known DDR modulators for greater sensitivity and reproducibility. In addition, we have characterized the most clinically relevant PARP inhibitors based on inhibition of PARylation and PARP1 trapping ability, in order to look for patterns of induced sensitivity to PARP inhibition in the presence of key DDR defects. We report here that clinically relevant PARP inhibitors can be functionally clustered into two unique classes, based on activity in the presence of base excision repair (BER) defects, and not on PARP1 trapping ability as was previously suggested. These results show that effectors of response to PARP inhibitors extend beyond the scope of HR perturbation and PARP trapping, and suggests that a better understanding of secondary targets may be critical for the optimal application of the numerous PARP inhibitors which are now being used in the clinic.

## RESULTS

### Clinically-relevant PARP inhibitors have varying degrees of specificity for PARP1 trapping and inhibition of PARylation

PARP inhibitors have traditionally been evaluated via qualitative or quantitative immunoassays measuring downstream PARylation in the setting of induced DNA damage. We adapted this format to 96-well microplates to better accommodate high-throughput quantification, and to more accurately parallel the methods used in short-term viability assays using the same compounds (**Fig 2A**). We tested a diverse and structurally unique collection of PARP inhibitors in these studies (**Fig 1**). As expected, all PARP inhibitors showed dose-dependent inhibition of PARylation in the setting of alkylation damage, with a nearly 1000-fold difference between the most potent inhibitor of PARylation, talazoparib, and the weakest tested, A-966492 (**Fig 2B**).

**Fig 1.**
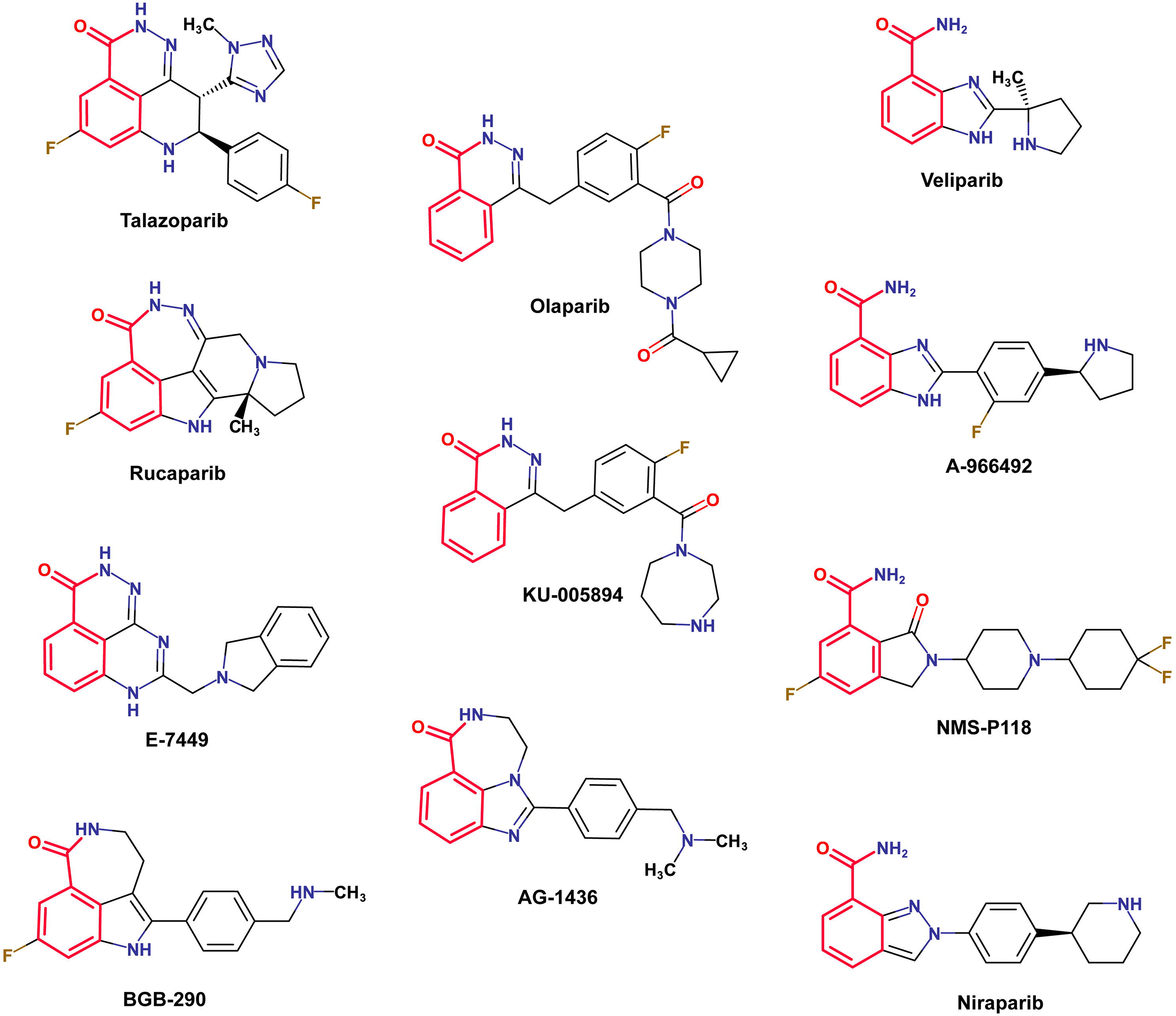
Chemical structures of selected PARPi. Highlighted in red is the 3-aminobenzamide - like structure which is thought to block PARP function via inhibition of NAD+ binding.

**Fig 2.**
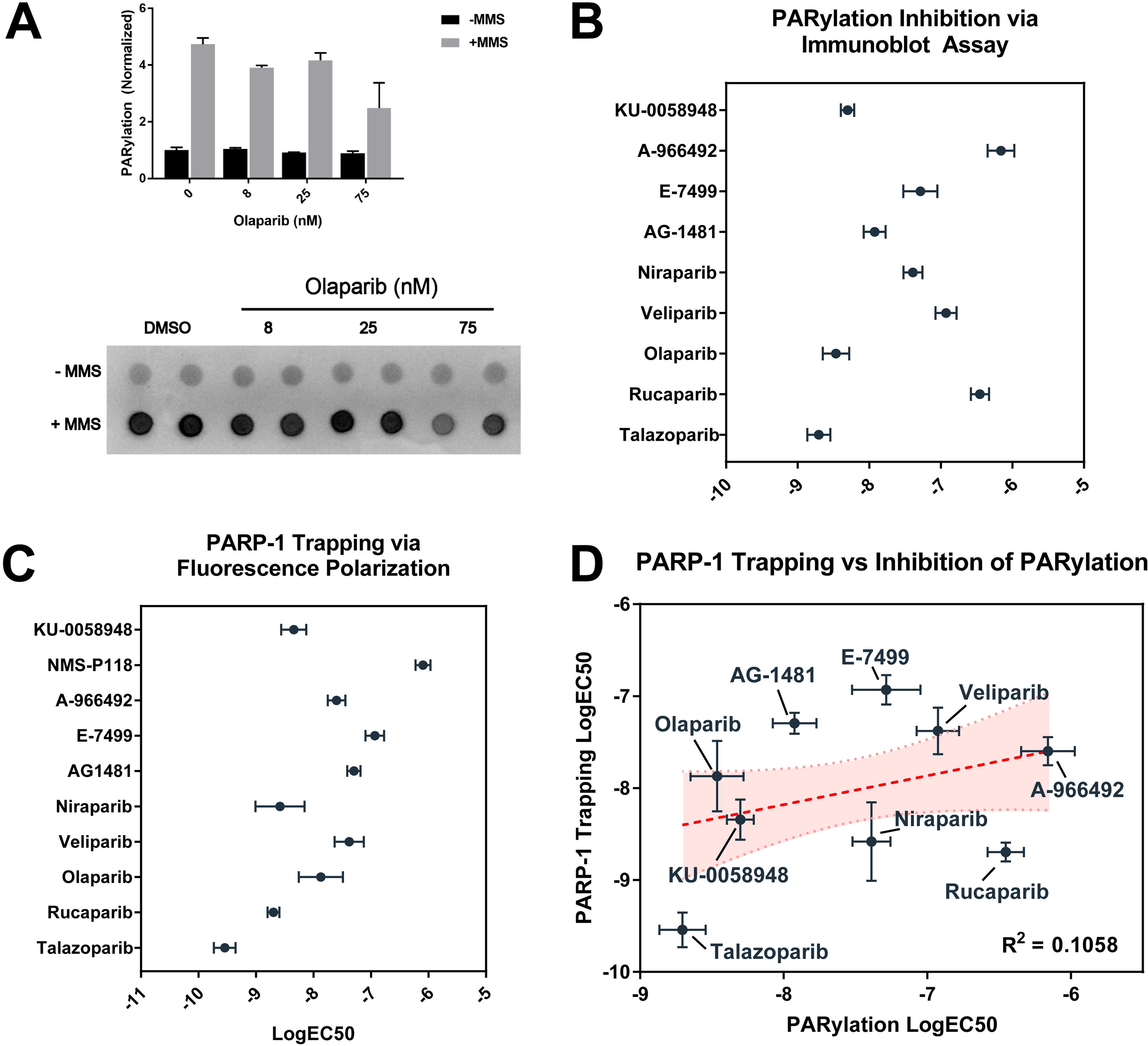
Biochemical characterization of PARPi shows limited correlation between inhibition of PARylation and PARP trapping. (A) Representative example of PARylation immunoassay in MCF10A cells in presence and absence of 0.01% MMS and increasing amounts of the PARP inhibitor olaparib. (B) Catalytic inhibition of PARylation EC50 as measured by PARylation immunoassay +/− SEM. (C) PARP1 trapping EC50 per agent as measured by fluorescence polarization assay +/− SEM. (D) Comparison of PARP1 trapping potency and catalytic inhibition of PARylation.

We then tested the same panel of inhibitors in a fluorescence polarization-based assay, which measures binding of PARP1 to a fluorescently-labeled DNA substrate in the presence and absence of PARP inhibition [12]. As expected, measured polarization of wells containing compounds reported to have strong PARP-trapping characteristics showed increased potency when compared to compounds such as veliparib, which have been reported to have limited trapping potency (**Figs S1A-B**). Similar to results from measured inhibition of PARylation, talazoparib was again found to be the most potent compound tested in the fluorescence polarization assay, with a measured IC50 approximately 10-fold lower than the next most potent compound (**Fig 2C**). Additional results were found to correlate well with previously published data [12, 19]. Notably, potency of PARP inhibitors as measured by PARylation immunoassay was not found to be significantly correlated with trapping potency as measured by PARP1 trapping assay (R^2^ = 0.1058, p > 0.05, Spearman r = 0.3), indicating that these two processes occur independent of one another (**Fig 2D**).

### Both PARP1 trapping potency and inhibition of PARylation fail to independently predict synthetic lethality in HR-deficient cells

As noted earlier, synthetic lethal interactions between PARP1 inhibition and HR-deficiencies are hypothesized to be markedly enhanced by trapping of PARP1 at sites of DNA damage [20]. In order to quantify growth inhibition across all tested PARP inhibitors, we performed short-term viability assays in isogenic HR-proficient and -deficient colorectal adenocarcinoma cell lines, DLD-1 and DLD-1 BRCA2−/−, respectively. Growth inhibition in both cell lines across the spectrum of PARP inhibitors was found to vary widely relative to the IC50s for PARylation and PARP1 trapping (**Fig 3A,B**). Growth inhibition in the HR-proficient DLD-1 cell line was found to be significantly correlated with both inhibition of PARylation (p = 0.006) and trapping potency (p < 0.0001) (**Fig 3A**). However, growth inhibition in HR-deficient DLD-1 BRCA2−/− cells was not found to correlate with inhibition of PARylation (p = 0.345) and only trended towards a significant correlation with trapping potency (p = 0.068) (**Fig 3B**). Interestingly, specific growth inhibition in HR-deficient cells relative to wild-type counterparts did not correlate with either inhibition of PARylation (p = 0.4384) or PARP1 trapping (p = 0.7213) (**Fig 3C**). Overall, these findings suggest that neither the inhibition of PARylation, nor the PARP trapping ability of PARP inhibitors independently predicts the magnitude of synthetic lethality of PARP inhibitors in HR-deficient cell lines.

**Fig 3.**
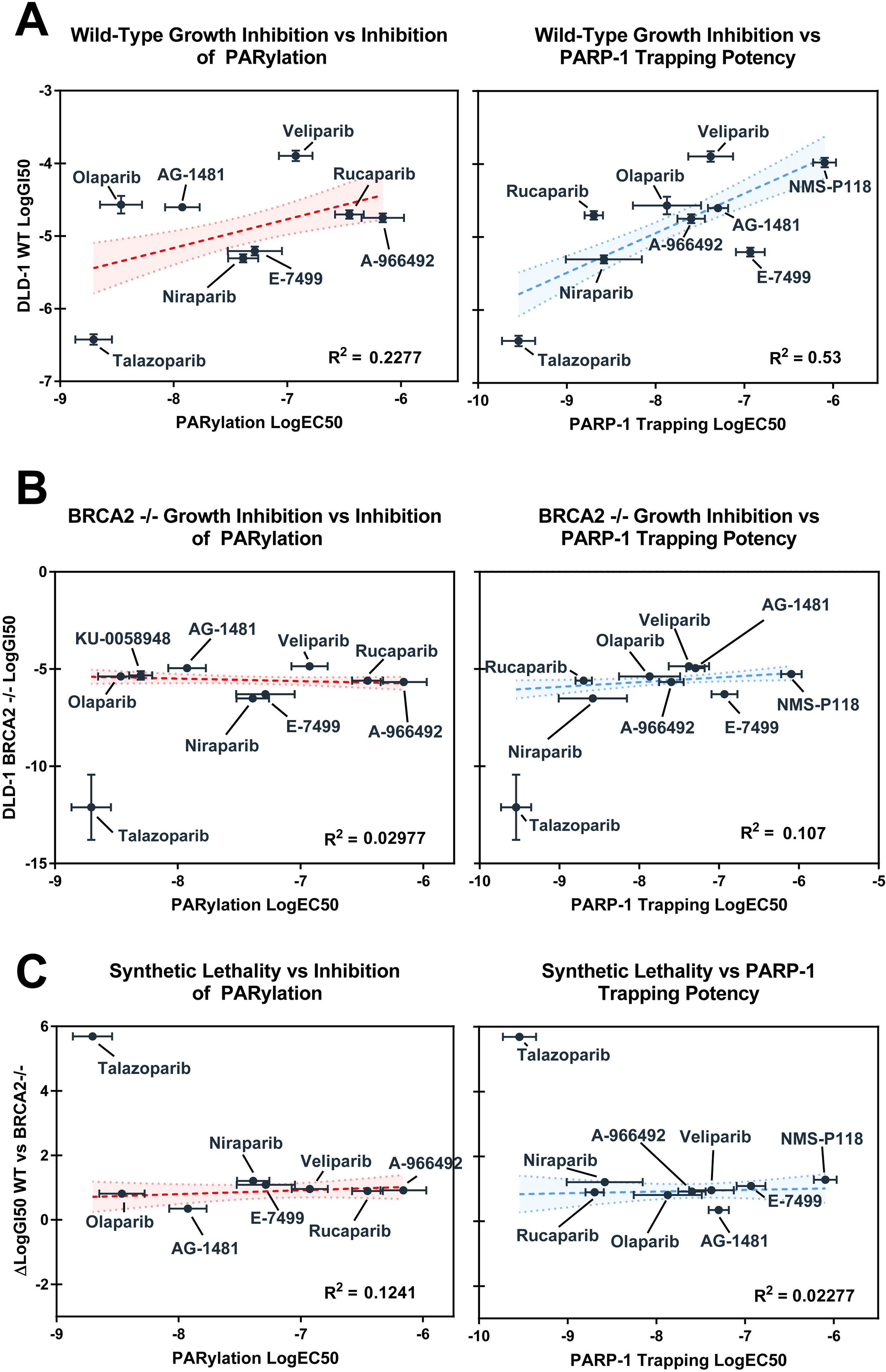
Synthetic lethality in HR-deficient cells shows no correlation with inhibition of PARylation nor PARP trapping potency alone. (A) Correlation between PARP1 trapping potency or inhibition of PARylation with growth inhibition in HR-proficient and -deficient DLD-1 cells. Correlation between growth inhibition in HR-proficient cells and inhibition of PARylation was significant (R^2^ = 0.2277; p = 0.006), as was correlation with PARP1 trapping potency (R^2^ = 0.53, p < 0.0001). (B) In DLD-1 BRCA2^−/−^ cells, there was no observed correlation between growth inhibition and inhibition of PARylation (R^2^ = 0.0298, p = 0.345), and correlation with PARP1 trapping only trended towards significance (R^2^ = 0.107, p = 0.068). (C) Synthetic lethality does not correlate with either strength of inhibition of PARylation or trapping potency.

### Targeted CRISPR/Cas9 screen reveals a novel classification of PARP inhibitors

Numerous prior studies have elucidated synthetic lethal interactions between PARP inhibition and specific DNA repair deficiencies (reviewed in [21]), while few have focused on possible differences between multiple structurally unique PARP inhibitors and DDR genes. We thus performed a targeted CRISPR/Cas9-based lentiviral screen using five structurally unique inhibitors from **Fig 1**, which were selected to represent the broad range of PARP trapping and PARylation activities that we observed in our earlier studies (see **Figs 4A,B**).

**Fig 4.**
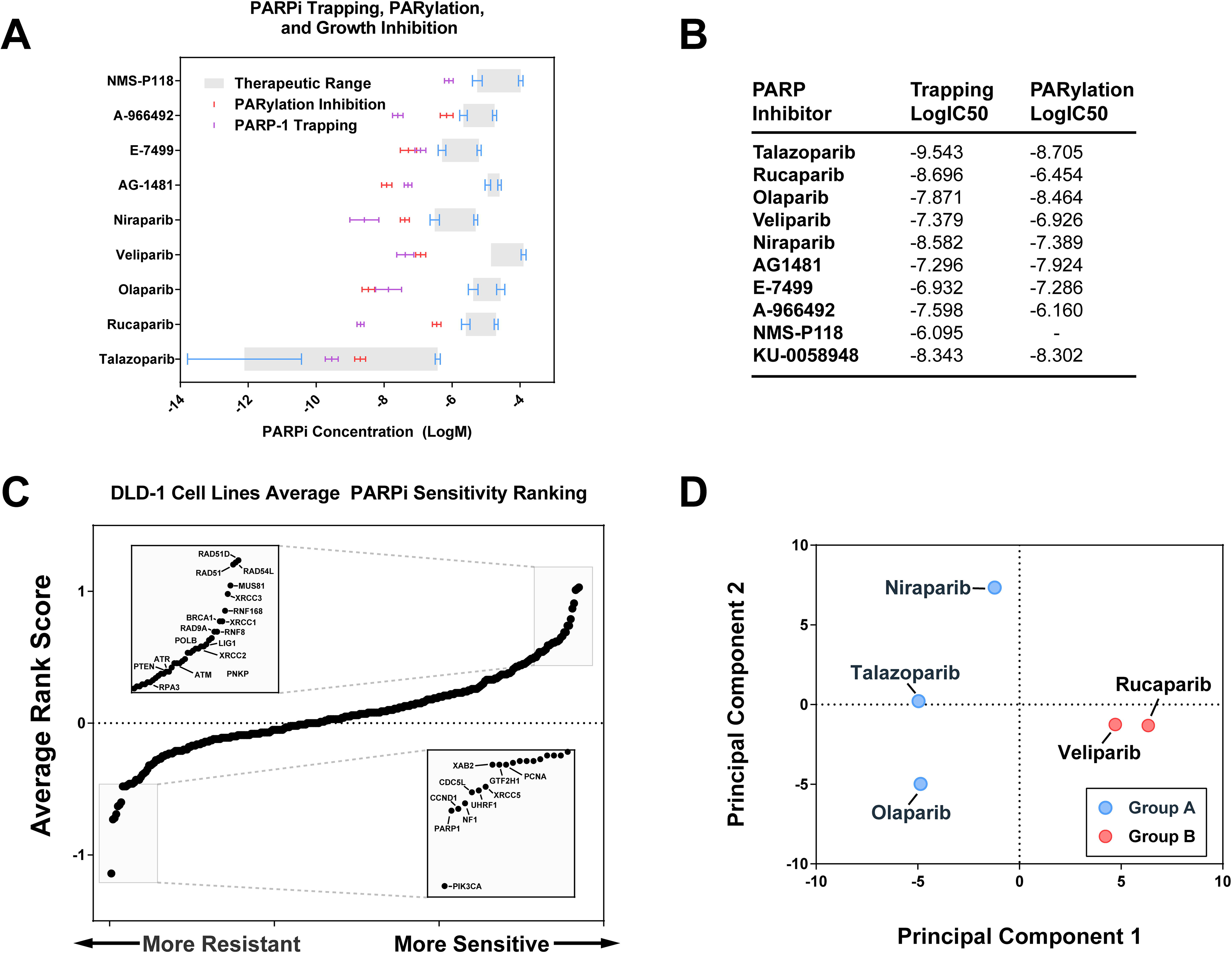
Targeted CRISPR screen reveals two unique functional groups of PARPi based on overall response to loss of DDR genes. (A) Visual representation of logIC50 values for trapping potency, inhibition of PARylation, and growth inhibition in HR-proficient and - deficient DLD-1 cells. (B) Summary of results from A. (C) Rank order average of tested inhibitors over entire screening set with single-gene knockouts conferring increased sensitivity or resistance highlighted at the extremes. (D) Principal components analysis of tested inhibitors reveals two distinct groups of inhibitors with talazoparib, olaparib, and niraparib making up Group A and veliparib, rucaparib making up Group B.

To evaluate the validity and sensitivity of our assay, analysis of all tested PARP inhibitors were combined in comparison to DMSO-treated control group, with the expectation that key proteins involved in homologous recombination would be among the most sensitizing findings. Among the top single-gene knockouts conferring sensitivity to all tested PARP inhibitors were RAD51, XRCC3, BRCA1, RNF8, ATM, ATR, and others (**Fig 4C**). Knockout of PARP1 was also shown to confer a general resistance to PARP inhibition as expected, though the size of this effect varied depending on the specific inhibitor in question. Individual PARP inhibitors were generally well-correlated with the average response to PARP inhibitors, with talazoparib being most similar to the average (R^2^ = 0.7307) and veliparib and rucaparib (R^2^ = 0.61, 0.614) being least correlated to the average response (**Fig S2**).

In order to look for trends in response to single-gene knockouts across multiple inhibitors, dimensionality reduction was performed, using response to each gene as input. Using these techniques, compounds showing similar responses across our targeted library should cluster closer together. Principal components analysis of inhibitors based on response to single-gene knockouts revealed two groupings of clinical PARP inhibitors, with Group A consisting of talazoparib, olaparib, and niraparib and Group B consisting of veliparib and rucaparib (**Fig 4D**). These data suggest a novel division of clinically relevant PARP inhibitors based entirely on functional classification in response to deficiencies in DNA repair, and does not appear to correlate with measured inhibition of PARylation or PARP1 trapping potency (**Fig 3A-C**).

### Group-specific targets reveal response to XRCC1, LIG3, and PARP1 knockout as key predictors of overall potency of PARP inhibitors

To better evaluate the defining characteristics between Group A and Group B inhibitors, we used publicly available gene ontology data to look for differences in effect of key DDR pathways. Although HR and Fanconi Anemia pathways showed the strongest sensitizing phenotype to both Group A and Group B inhibitors, differences between the two groups were best exemplified by differences in sensitization to key proteins in both base excision repair (BER) and mismatch repair (MMR) pathways (**Fig 5A**). Within BER, increased resistance to PARPi in the presence of PARP1 knockout and increased sensitivity to PARPis upon loss of LIG3 and XRCC1 were the most defining characteristics of Group A inhibitors relative to others (**Fig 5B**). Increased sensitivity to POLE4 was also noted among Group A inhibitors followed by differential sensitivities to FEN1, LIG1, and PARP3 approaching significance (**Fig 5B**).

**Fig 5.**
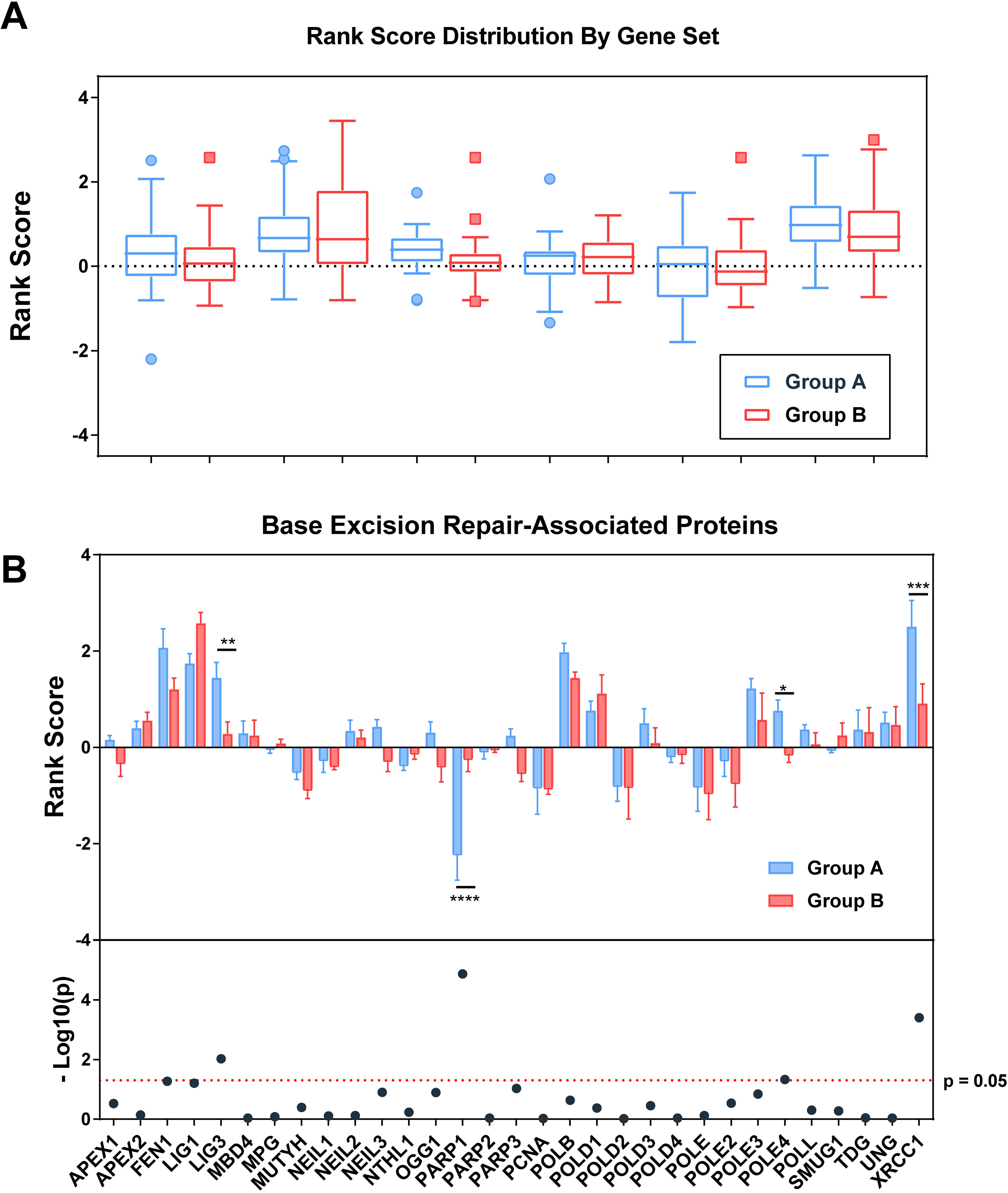
Inter-group variation in response to PARPi shows greatest difference in genes associated with BER. (A) Comparison of group-averaged rank score by associated pathway. Higher rank scores are associated with increased sensitivity to loss of function of proteins within each reported pathway. A single gene may appear in more than one pathway. (B) Per-gene rank scoring by PARPi group reveals significant differences in PARP1, LIG3, XRCC1, and POLE4 response to inhibitors (p<0.0001, p=0.0094, 0.0004, and 0.047 via student’s t-test). Additional genes involved in base excision repair approaching significance include FEN1, LIG1, and PARP3 (p=0.054, 0.062, 0.093).

Findings from the initial screen were confirmed first by testing selected sgRNAs from the original library by 96h short-term viability assay and then by pooled siRNA experiments to measure the effect of knockdown, rather than knockout, of each gene in the presence of PARPi. Short-term viability assays were also performed using U2-OS cells to show effects carry across unrelated cell lines, independent of tissue of origin. Both individual sgRNA experiments, as well as siRNA experiments, largely recapitulated the results seen by pooled CRISPR/Cas9 screening (**Figs 6A-C, Fig S3**). XRCC1 and LIG3 knockouts and knockdowns show increased sensitivity to Group A inhibitors that are far less pronounced or absent in Group B across all assays. These findings confirm the results from our targeted CRISPR/Cas9 screen, showing that loss of function of XRCC1 and LIG3 confer increased sensitivity to some, but not all PARP inhibitors. Interestingly, sensitization to PARP inhibition has been shown previously in the setting of XRCC1 deficiency, however this study was limited only to the Group A inhibitors, talazoparib and olaparib [22]. Additionally, the degree of sensitization in the setting of loss of either XRCC1 and LIG3 appears to correlate with the overall PARP1-dependence of toxicity, and may provide critical insight into better understanding the therapeutic effects of PARP inhibition in the setting of such deficiencies.

**Fig 6.**
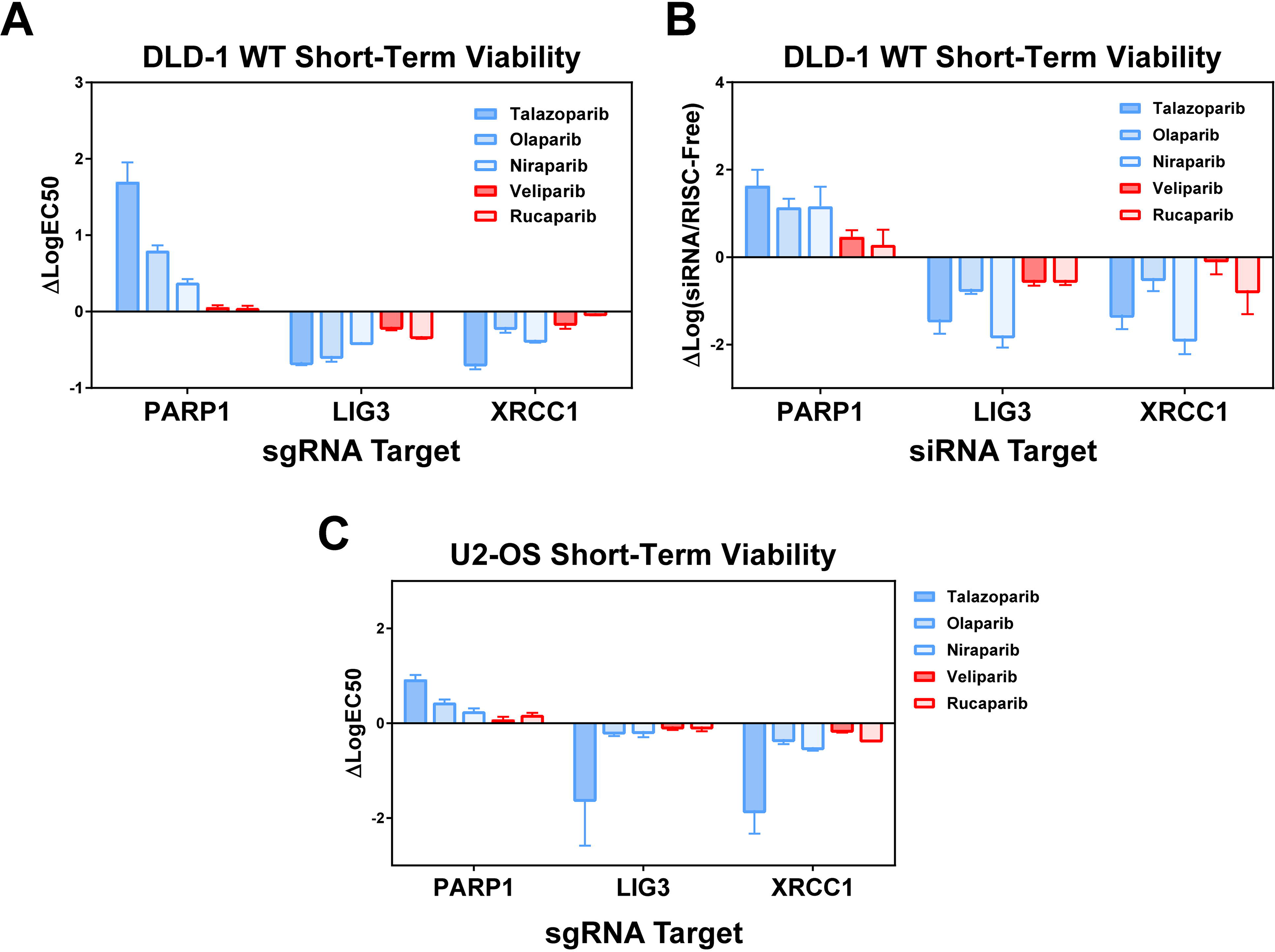
Loss of XRCC1 and LIG3 increase sensitivity to Group A inhibitors with limited effect to Group B. (A) Short-term viability assays reveal specific response to PARP1, LIG3, and XRCC1 seen from CRISPR/Cas9 screen. Strong resistance in presence of PARP1 knockout also associated with increased sensitivity in presence of XRCC1, LIG3 knockout. (B) Pooled siRNA knockdown of each of the reported genes shows similar phenotype to CRISPR/Cas9 lentiviral knockout; again showing increased sensitivity to talazoparib, olaparib, and niraparib in the presence of XRCC1/LIG3 disruption. (C) Short-term viability assays in U2-OS cell line shows similar phenotype with lentiviral CRISPR/Cas9 knockout of reported genes.

### In *silico* analysis reveals similar clustering of tested PARP inhibitors and association with PARP1/XRCC1/LIG3 loss-of-function

To assess our functional genetic screening methods in comparison to alternative datasets, we examined publicly available datasets from the DepMap project comparing gene essentiality and drug sensitivity across hundreds of human cell lines. Interestingly, principal components analysis of the relative sensitivities of each of the five examined PARP inhibitors tested across over 400 cell lines reveals clusters coinciding with those found via our functional genetic screen, with higher degrees of correlation between talazoparib, niraparib, and olaparib than with rucaparib and veliparib (**Fig 7A**). We next performed hierarchical clustering to identify groupings of cell lines with similar sensitivity patterns to the tested inhibitors to examine qualities relating to selective sensitivity in talazoparib, niraparib, and olaparib (**Fig 7B**). Notably, cell lines with relative sensitivity to PARP inhibition were split between two groups - the pan-sensitive group identified as Cluster 1 and the selectively sensitive group identified as Cluster 3. Cell lines within Cluster 3 are defined by moderate to high sensitivity to talazoparib, niraparib, and olaparib and mid to low sensitivity to rucaparib and veliparib (**Fig 7C**).

**Fig 7.**
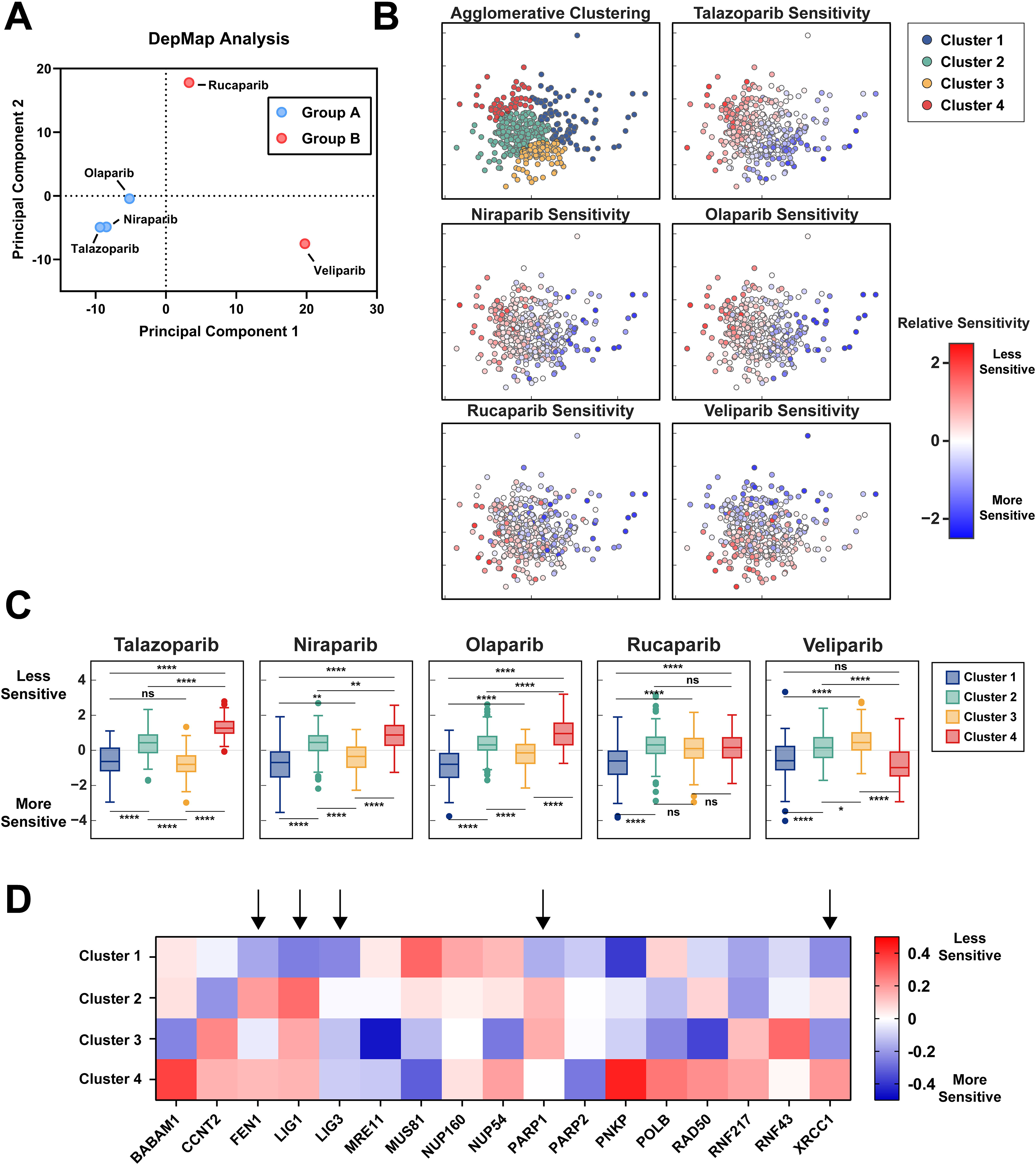
Gene essentiality and drug sensitivity studies from DepMap dataset confirm functional clustering of PARPi and dependence on BER pathway. (A) Principal components analysis of DepMap sensitivities to tested PARP inhibition shows clustering similar to those seen via PCA of our functional genetic screen with talazoparib, olaparib, and niraparib having similar effects across cell lines in comparison to rucaparib and veliparib. (B) Principal components analysis and agglomerative clustering of cell lines in response to PARP inhibition reveals 4 distinct clusters. (C) Clustered sensitivity to PARP inhibition showed compound-specific responses, particularly in Cluster 3, which shows equal sensitivity to the pan-sensitive Cluster 1 in talazoparib, similar sensitivity to Cluster 1 in niraparib and olaparib, but relative resistance in comparison to Cluster 1 in rucaparib and veliparib. (D) Selection of genes showing differences in essentiality across clusters. Columns denoted by arrows correspond to genes found to differentially affect response to PARP inhibition between Group A and Group B inhibitors.

We next performed an analysis of gene essentiality within each cluster to measure what factors correlate with pan-sensitivity rather than selective sensitivity. Analysis of relative sensitivity to loss of selected genes identified within our functional screen as well as genes having a significant cluster-dependent effect via ANOVA analysis (**Fig 7D**). Cell lines exhibiting pan-sensitivity to PARP inhibition in Cluster 1 are more sensitive to loss of FEN1, LIG1, PARP1, and PARP2, but not LIG3 or XRCC1 in comparison to cell lines showing selective sensitivity in Cluster 3. Such differences between pan-sensitive and selectively sensitive cell lines may provide insight into the differing mechanism resulting in cytotoxicity in the presence of PARP inhibition.

## DISCUSSION

While the exact mechanistic basis for synthetic lethal interactions with PARP inhibition in the setting of BRCA1/2 mutations and HR deficiencies remains controversial, our data clearly demonstrate that neither trapping potency nor strength of inhibition of PARylation fully explain the response to such inhibitors. These findings are in agreement with recent biochemical studies suggesting that inhibitors of PARP1 fit into three major classifications based on allosteric effects of PARPi binding as well as retention at sites of DNA damage [23]. Similarly, our unbiased analysis of over 280 genes known to be involved in DNA damage repair and response found unique groupings of PARP inhibitors which do not correlate solely with either the ability to inhibit downstream PARylation by PARP1 or the trapping of PARP1 to sites of damage based on widely-used biochemical assays. Across the PARP inhibitors tested in our analysis, we do not observe any correlation between synthetic lethality in the context of HR defects and strength of PARP1 trapping or inhibition of PARylation. Indeed, PARP trapping has been associated with increased toxicity in both normal tissue as well as within tumors, likely resulting in side effects seen in clinical trials such as complete bone marrow failure and other cytopenias [24]. Such findings make appropriate classification of inhibitors for use in patient populations ever more relevant, as the use of PARP inhibitors in clinic becomes increasingly common.

Within our screen, we see strong sensitization to all PARP inhibitors through knockout of key components of HR (RAD51, BRCA1, BRCA2, etc.), however only three of our tested inhibitors respond to loss of function of proteins immediately downstream of PARP1 in BER. Interestingly, loss of XRCC1 and LIG3 was found to be most toxic to cells concurrently treated with inhibitors that are dependent on PARP1 for sensitization (Group A PARP inhibitors). We hypothesize that this observation may be due to one or more of the following mechanisms. i) PARP1-independent inhibitors may be maximally disrupting downstream BER through disruption of PARP1 signaling at lethal doses, so further loss of function does not alter response to inhibition. ii) Loss of XRCC1 and LIG3 results in hyperactivation of PARP1 as has been shown previously, and is therefore increasing opportunities for PARP1-dependent toxicity [25]. iii) Loss of XRCC1 and LIG3 results in unrepaired lesions of the DNA, which may be preferentially targeted by PARP1-dependent inhibitors. Differential sensitivity to loss of function of PARP1, PARP2, LIG1, and FEN1 as seen in our DepMap essentiality analysis indicates that all PARP inhibitors may be equally effective in targeting cells dependent on BER function, however talazoparib, niraparib, and olaparib may have extended functionality outside of this scope. Additional work is necessary to tease apart such mechanisms and further evaluate the utility of various classes of PARP inhibitors in specific clinical settings.

Although there are over 250 active clinical trials testing PARP inhibitors in cancer at the time of this writing, there is little information regarding appropriate selection of PARP inhibitor therapy and utilization of PARP inhibitors in patients who have failed to respond to one or more of such inhibitors. Likewise, no head-to-head clinical trials comparing PARP inhibitors have been completed to date, making selection of PARP inhibitor treatment in the clinical setting difficult. Neither PARP trapping nor catalytic inhibition of PARylation appear to explain the efficacy of PARP inhibition in the treatment of cancers with DNA repair deficiencies. Our results indicate that the efficacy of PARP inhibitors may hinge on some combination of PARP trapping and inhibition of downstream targeting of PARP1, with a handful of inhibitors, talazoparib > niraparib > olaparib, being far more dependent on the presence of PARP1 than others. Clinical trials are necessary to determine the utility of PARP1-independent inhibitors in the setting of limited PARP1 expression. Additionally, patients with mutations in XRCC1 and LIG3 may benefit from treatment with talazoparib, olaparib, or niraparib over treatment with PARP1-independent inhibitors. Further studies are necessary to determine how these results may affect response to treatment in patients, and whether our findings may translate into a clinical setting. Overall, our results highlight an exciting technique in functional analysis of PARP inhibition via CRISPR/Cas9 screening to define genetic dependencies, and show the importance of functional BER in the setting of select PARP inhibitors.

## MATERIALS AND METHODS

### Cell lines and reagents

Colorectal adenocarcinoma cell lines DLD-1 and DLD-1 BRCA2 −/− were used and maintained in RPMI medium with 10% fetal bovine serum (FBS; Gibco) at 37°C with 5% CO_2_. The DLD-1 BRCA2^−/−^ cell line has an engineered exonic deletion as described [26]. HEK293FT (ThermoFisher) are human embryonal kidney cells maintained in Dulbecco’s Modified Eagle’s Medium, high glucose (DMEM; Thermo Scientific/Gibco) supplemented with 10% FBS. MCF10A cells are normal human mammary cells and were maintained in DMEM/F12 media (Gibco, #11330032) supplemented with 5% horse serum, 10 ng/ml epidermal growth factor, 0.5 mg/ml hydrocortisone, 100 ng/ml cholera toxin, and 10 ug/ml insulin. PARP inhibitors tested for the purposes of this study were obtained from the following vendors: Talazopoarib (Selleckchem; #S7048), olaparib (Selleckchem; #S1060), rucaparib (Selleckchem; #S1098), niraparib (Selleckchem; #S2741), veliparib (Selleckchem; #S1004), A-966492 (Selleckchem; #S2197), KU-0058948 (Axon Medchem; #2001), NMS-P118 (Selleckchem; #S8363), E-7449 (Selleckchem; #S8419), AG-014699 (Axon Medchem; #1529), BGB-290 (BeiGene).

### PARylation Immunoblot

To measure PARylation inhibition, MCF10A cells were plated on 96-well microplates (Greiner) at a density of 20k cells/well 24h prior to treatment with methyl-methanesulfonate (MMS; Sigma) and indicated PARP inhibitors. After 24h in culture, media from the plates was aspirated and a fresh 75 μl of pre-warmed media was added to each well. To this, 25 μl of media containing either 0.01% MMS, PARP inhibitors, or a combination of the two were added to each well. Cells were incubated for 30m in normal culture conditions. Following the 30m culture, media was aspirated and cells were rinsed once with PBS. Cell cultures were lysed with RIPA lysis buffer for 30m at 4°C with occasional agitation. Lysates were spotted on nitrocellulose membrane (BioRad) and allowed to dry at room temperature for 1h. Blocking was performed in TBS-T with 5% BSA (Gold Biotechnology) for 1h at room temperature, followed by overnight incubation with anti-PAR antibody (Trevigen, #4336-BPC-100) at 4°C. After primary incubation, three 10-minute washes with TBS-T were performed, followed by 1h incubation with HRP anti-rabbit conjugated secondary antibody (ThermoFisher; #31462) at room temperature under constant agitation. Images obtained on ChemiDoc (BioRad) following addition of Clarity Western ECL substrate (BioRad). Image quantification was done using ImageJ imaging software and normalized to no-MMS and no-PARPi control [27]. Curve fitting and data analysis performed using Graphpad Prism (Graphpad Software).

### PARP1 Trapping Assay

Preparation of PARP1 dsDNA substrate was performed as previously described [28]. Briefly, single-stranded oligonucleotides were hybridized by combining in equimolar ratio of the following sequences:

5′-AlexaFluor488-ACCCTGCTGTGGGCdUGGAGAACAAGGTGAT
ATCACCTTGTTCTCCAGCCCACAGCAGGGT

This mixture was then heated to 95 °C for 5m and slowly cooled to room temperature at 5 °C/min. Hybridized oligonucleotide was then incubated with APE1 and UDG (NEB) at 37°C for 1h to create a single strand break recognized by the PARP1 enzyme. To measure inhibition of release of DNA substrate from PARP1 enzyme, 30 nM GST-Tagged PARP1 protein (BPS Biosciences) was incubated with 1 nM DNA substrate and varying amounts of PARPi or DMSO for 1h in reaction buffer containing 50 mM Tris (pH 8.0), 4 mM MgCl_2_, 10 mM NaCl, and 50 ng/ml BSA in water at RT. After 1 hour, fluorescence polarization readings were recorded using a Cytation 3 (Biotek) multi-mode imager with fluorescence polarization filter prior to adding 1mM NAD+ and every 5 minutes after. Curve-fitting and statistical analysis was performed using Graphpad Prism (Graphpad Software). The concentration of PARP1 to fluorescent dsDNA substrate was first titrated to optimize detection of polarization via automated plate reader in 96-well half volume microplates (**Fig S1**). To measure trapping efficiency of various PARP inhibitors, purified PARP1 protein, DNA substrate, and varying concentrations of PARP inhibitors were incubated for 1h at room temperature to ensure saturated binding capacity. After the incubation, NAD+ was added to the reaction to initiate release of DNA from PARP1, and polarization measurements were recorded in 5-minute intervals for 120 minutes. Importantly, controls lacking NAD+, PARP1 protein, and DNA substrate were included for normalization.

### CRISPR/Cas9 Screening and Analysis

A CRISPR/Cas9 DDR targeted library was assembled using available gene ontology databases and lists of genes involved in DNA damage repair and response. The top 10 suggested sgRNAs targeting each gene were selected from the http://www.genome-engineering.org/ website and supplemented with non-targeting control sgRNAs [29]. These oligos were assembled into the LentiCRISPRv2 lentivirus backbone as described in the original protocols [30, 31]. Viral production was carried out in HEK293FT cells by equimolar co-transfection of LentiCRISPRv2 library, psPAX2, and pCMV-VSV-G using Lipofectamine 2000 (Invitrogen; #11668027). lentiCRISPR v2 was a gift from Feng Zhang (Addgene plasmid # 52961), psPAX2 was a gift from Didier Trono (Addgene plasmid # 12260), and pCMV-VSV-G was a gift from Bob Weinberg (Addgene plasmid # 8454) [32]. Viral titer was assessed upon collection and concentration of lentiviral supernatant with Lenti-X Concentrator (Takara Biotech; #631231). Appropriate final concentrations were chosen to maintain MOI < 0.3 to reduce probability of coinfection with two or more sgRNA sequences. For screening, DLD-1 cells were transduced in 8 μg/ml Polybrene (Sigma-Aldrich) with the multiplexed CRISPR/Cas9 library containing 10 unique sgRNAs targeting 284 different genes involved or implicated in DDR-associated pathways along with one thousand non-targeting sgRNA controls. Cells were then selected with Puromycin (InvivoGen; #ant-pr-1) for 3 days following transduction, ensuring a MOI < 0.3 to prevent multiple sgRNA integrations per cell. After initial selection, cells were split into six treatment groups and treated with appropriate PARPi at calculated GI_30_ or DMSO as indicated to assess effects on both sensitivity and resistance to tested inhibitors. Samples were taken at Day 0 as well as every 2 days to ensure logarithmic growth while maintaining a high sample size.

Preparation and sequencing of samples was conducted using dual-indexed paired-end sequencing on MiSeq System (Illumina) using a 2×150 protocol. Library preparation was conducted two independent primer sets. Primers used in the first reaction amplify the targeted sgRNA region of the integrated vector and primers used in the second reaction allow for indexing and multiplexing during sequencing. Additional spacer sequences of 0-2 bases were inserted between the adapter and sequence-specific portions of the sequencing primers to increase library diversity during sequencing.

Analysis was performed using a rank scoring algorithm similar to one previously described [33]. sgRNAs were extracted from sequencing reads, counted, and normalized to total sample size and non-targeting control abundance. A rank score was calculated for each gene represented in the targeted library by comparing the abundance of each sgRNA to its representation in the targeting library sample. Each screen was done in duplicate and samples were prepared from multiple time points in each treatment group to reduce sampling error.

### Short-Term Viability Assays

Short-term viability assays validating individual sgRNA results were performed by first transducing cells in 6-well plates with lentivirus, selecting with Puromycin for 48h, then plating into 96-well plates for 24h prior to adding appropriate concentrations of PARPi. 96h after the initiation of treatment, media was aspirated, cells were washed once with PBS and were then fixed using 4% formaldehyde (Sigma-Aldrich) in PBS solution for 15m at room temperature. Cells were then stained with Hoechst 33342 (Sigma-Aldrich; #B2261) for 45 minutes prior to imaging using the Cytation 3 multi-mode imager as described previously [34]. Cell counting was performed using a pipeline created in CellProfiler image analysis software which stitches images by well and identifies the number of cell nuclei per well by fluorescence staining [35]. Graphing and data analysis was performed using Graphpad Prism (Graphpad Software). Assays utilizing pooled siRNA (Horizon; ON TARGETplus siRNA) were conducted by first transfecting with RNAiMAX (Invitrogen; #13778100) 72h prior to exposure to individual PARP inhibitors to ensure maximum knockdown at initial treatment.

### In Silico Analysis of Drug Sensitivities and Gene Essentiality

Gene essentiality data, drug sensitivity data, and accompanying cell line information was obtained via the DepMap Data Portal (https://depmap.org/portal/download/) with preprocessing steps as described in accompanying manuscripts [36–39]. Additional data processing, dimensionality reduction, and plotting was done via Scikit-Learn [40].

## ACKNOWLEDGEMENT

This work was supported in part by the US National Institutes of Health (NIH) grant R01CA215453.

