## Supplemental Figures for "CRISPR screening identifies novel PARP inhibitor classification based on distinct base excision repair pathway dependencies"

**Supporting Information**

S1

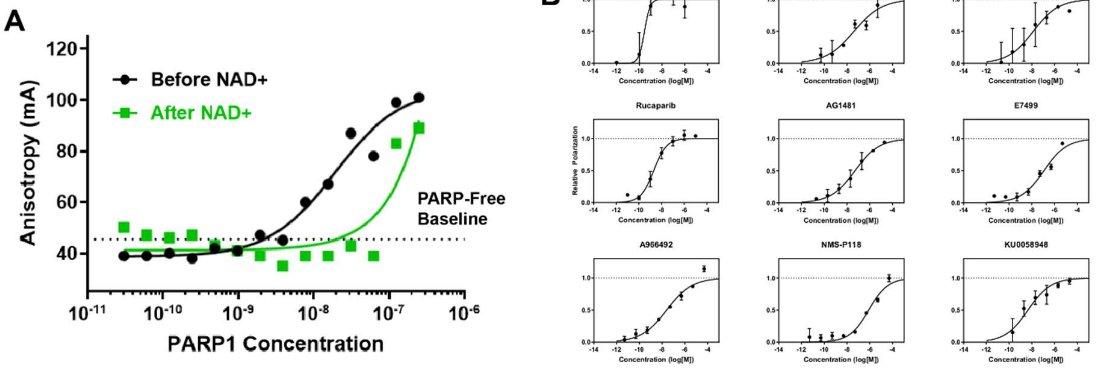


**Figure S1. PARP trapping potency measured across tested inhibitors via fluorescence polarization. A.** Measured anisotropy before and after addition of NAD+ to PARP1 and fluorescently labelled DNA substrate solution. PARP1 concentration chosen to maximize sensitivity to bound and unbound DNA substrate after the addition of NAD+. **B.** Dose-response curves for each of the tested PARP inhibitors for measurement of PARP-trapping properties by fluorescence polarization.

S2

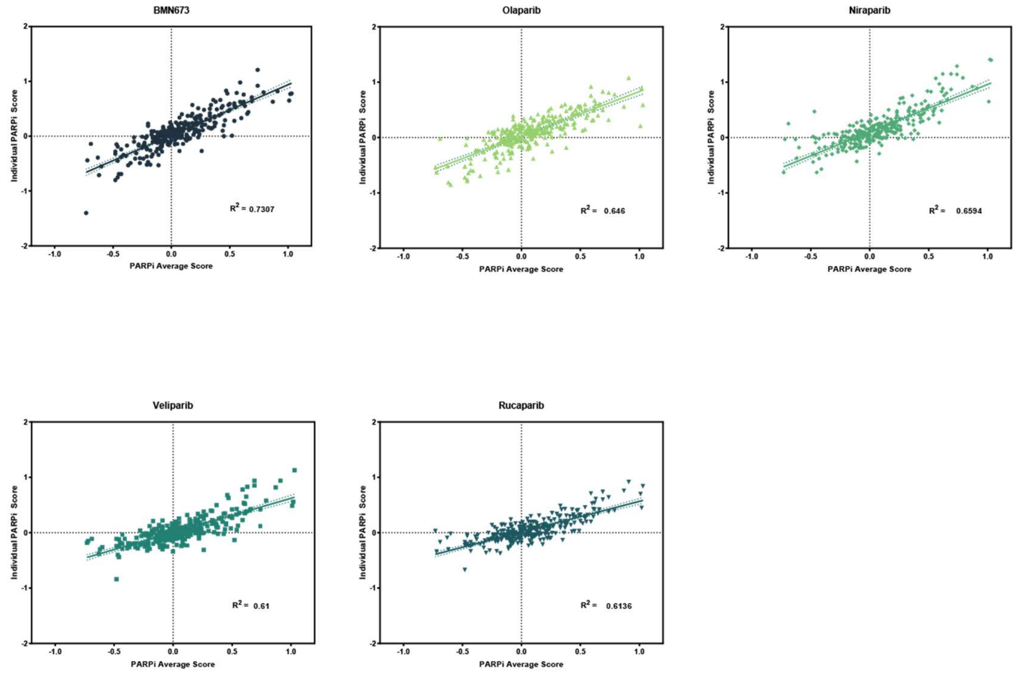

**Figure S2. Intra-class gene essentiality correlation between each inhibitor and the average across all tested PARPi.** Relative correlation between each individual PARP inhibitor and the average inhibitor. Each data point represents a single gene tested in the CRISPR/Cas9 screening library and the axes represent individual rank scores for each gene. Perfect correlation between each inhibitor and the average inhibitor would be represented by a straight line with $R^2 = 1$.

**S3**

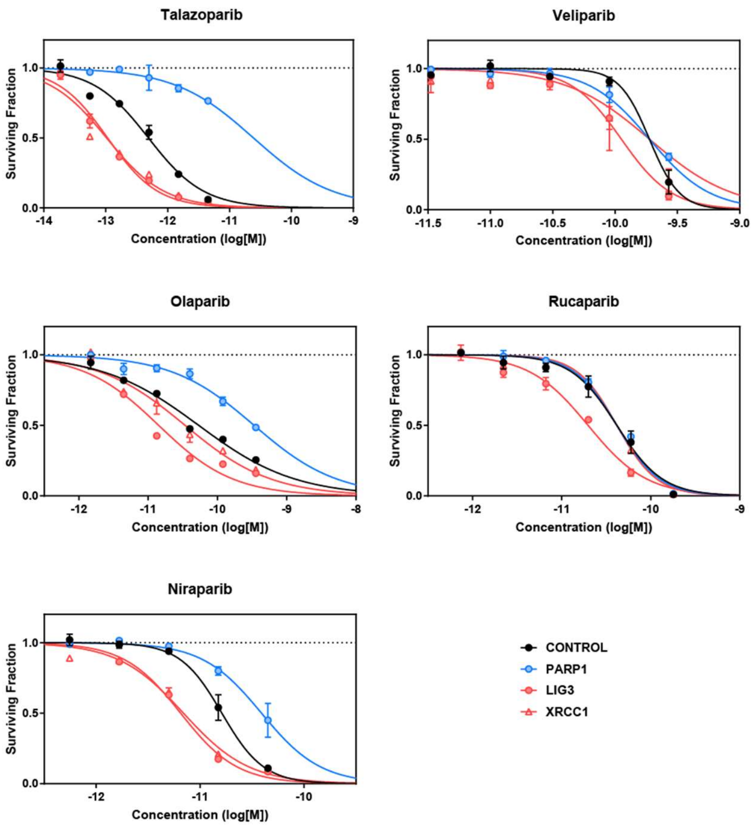


**Figure S3. Short term viability assays in DLD-1 cells and each knockout gene.** Dose-response curves for DLD-1 cells following lentiviral CRISPR/Cas9 knockout of each gene as indicated. Effect on IC50 for each of these graphs is quantified in Fig 6A.
